# Interprotomer Crosstalk in Mosaic Viral Glycoprotein Trimers Provides Insight into Polyvalent Immunogen Co-assembly

**DOI:** 10.1101/2025.04.22.649918

**Authors:** Chengbo Chen, Klaus N. Lovendahl, Julie M. Overbaugh, Kelly K. Lee

**Affiliations:** Department of Medicinal Chemistry, University of Washington, Seattle, WA 98195, USA; Biological Physics Structure and Design Graduate Program, University of Washington, Seattle, WA 98195, USA; Human Biology Division, Fred Hutchinson Cancer Center, Seattle, WA 98109, USA

## Abstract

SARS-CoV-2 variants have demonstrated the ability to evade immune responses, leading to waves of infection throughout the pandemic. In response, bivalent mRNA vaccines, encoding the original Wuhan-Hu-1 and emerging variants, were developed to display both spike antigens. To date, it has not been determined whether co-transfection and co-translation of different SARS-CoV-2 variants results in co-assembly of mosaic heterotrimer antigens and how this may affect trimer stability, dynamics, and antigenicity. Understanding whether such mosaic heterotrimers can form and their implications for antigen structure can provide important information to guide future polyvalent vaccine design where multiple variants of an antigen are co-formulated. To investigate this, we purified mosaic spike assemblies of both genetically close (Omicron BA.2 and XBB) and distant (Omicron BA.2 and Wuhan-Hu-1 G614) strains. We found that the stability and integrity of mosaic spike trimers were maintained without misfolding or aggregation. Glycosylation profiles likewise were preserved relative to the homotrimer counterparts. Hydrogen/deuterium-exchange mass spectrometry and biolayer-interferometry were used to investigate the mosaic spike dynamics and any impact on epitope presentation and receptor binding. The Omicron-XBB heterotrimer, sharing a common fusion subunit sequence, retained protomer-specific dynamics similar to the corresponding homotrimers in antigenically important regions. The Omicron-G614 heterotrimer, co-assembling from protomers of divergent fusion subunit sequences, likewise showed overall similar dynamics and conformations in the receptor-binding subunit compared to the homotrimers. However, the incorporation of the Wuhan-Hu-1 G614 protomer led to a stabilizing effect on the relatively unstable Omicron fusion subunit in the heterotrimer. These findings reveal structural dynamic crosstalk in mosaic trimers, suggesting a potential for enhanced immunogen display and important considerations to be aware of in the use of polyvalent nucleic acid vaccines.

**Author summary:** Here we investigated the possibility of immunogen co-assembly from bivalent nucleic acid vaccines with a focus on probing potential impacts on antigen stability and epitope display. We purified mosaic heterotrimers composed of Omicron BA.2 and Wuhan-Hu-1 G614 protomers as well as those composed of Omicron BA.2 and XBB protomers. Both mosaic heterotrimers maintained their structural integrity, morphology, and N-glycosylation profiles. Using hydrogen/deuterium-exchange mass spectrometry, we demonstrated that both types of heterotrimers preserved strain-specific dynamics in critical antigenic regions, comparable to their homotrimer counterparts. Crosstalk between fusion subunits of Omicron BA.2 and Wuhan-Hu-1 G614 revealed a stabilizing effect of the G614 protomer on the inherently less stable fusion subunit in Omicron. Our findings provide insight into the structural dynamic profiles of co-expressed immunogen display, highlighting important considerations for polyvalent formulations that are being pursued to provide broad protection against highly variable pathogens.

## Introduction

SARS-CoV-2 has been circulating in the human population since 2019, undergoing rapid evolution and diversification under selective pressure from immune responses to infection and vaccination.(1, 2) Much of the resulting variation occurs in the viral Spike (S) glycoprotein, which mediates entry into host cells by attaching to Angiotensin-Converting Enzyme 2 (ACE2) receptor then carrying out the process of virus-host membrane fusion.(3–5) Understanding the effect of sequence variations in S is important for studies of virus adaptation in response to host immunization and development of effective vaccines.

The D614G mutation was one of the first significant mutations to emerge in S. Located at the interface of receptor binding (S1) and fusion (S2) subunits, this residue substitution was proposed to increase the stability of S and infectivity of the virus.(6) Structural studies using an engineered pre-fusion S ectodomain construct showed that D614G exhibits enhanced contacts at the interface of S1 and S2 subunits and a bias towards adopting a receptor binding domain (RBD) “up” configuration compared to the original Wuhan-Hu-1 WT (Hu-1) S.(7–13) Most mutations in S occur in the RBD, leading to increased affinity for human ACE2 and escape from neutralizing antibody (nAb) pressure.(14, 15) Indeed most nAb elicited after natural infections target the RBD on the S1 subunit where they block ACE2 receptor binding.(14–16) Notable hotspot mutations in the RBD, such as N501Y, E484K/A, and K417N/T, combined with D614G mutation, have led to peaks in infection waves associated with variants of concern (VOCs), before major vaccination administrated to the human population was initiated in 2021.(1, 17, 18)

Both antigenic drift and antigenic shift during SARS-CoV-2’s wide-spread propagation led to improve viral fitness. In late 2021, the Omicron variant emerged, bearing extensive mutations in the RBD and throughout the spike assembly, with enhanced ability to evade immune responses and increased transmissibility.(19, 20) Omicron spread rapidly worldwide and diversified into numerous subvariants such as BA.1, BA.2, and BA.5, and their offspring subvariants. XBB, which emerged later, has been suggested to be a recombinant strain following coinfection of a shared host cell by separate strains.(21–23) This recombinant nature allows XBB to have the N-terminal homologous amino acid sequence to variant BJ.1 up to position 459, and the C-terminal sequence from subvariant BM.1. XBB thus inherited structural features from both subvariants that confer enhancements in human ACE2 binding and immune evasion.(24, 25)

Structure-based vaccine design featuring the display of entire Hu-1 S or RBD antigens have shown protections in preventing severe infections.(26–31) In particular, the efficacy of mRNA vaccines encoding modified S proteins was demonstrated in the COVID-19 pandemic.(32, 33) The two most prominent mRNA vaccines developed for COVID-19, Pfizer-BioNTech’s BNT162b2 (26) and Moderna’s mRNA-1273 (29) vaccines, encoding full-length, two-proline-stabilized Hu-1 S (S-2P), demonstrated high efficacy on a global scale by leveraging the human body’s cellular machinery to produce antigens and elicit an immune response.(34–36)

The severe pandemic situation and rapid evolution of circulating virus raised the demand for vaccines with high efficacy that could provide broad-spectrum protection against co-existing and newly emerging variants. In response to the emergence of the Omicron VOC’s circulation, a special bivalent mRNA booster vaccine was developed and deployed in fall 2022. This bivalent vaccination contained a 1:1 ratio mix of mRNA-LNPs encoding ancestral Hu-1 and Omicron variants and delivered to human cells with the aim of producing both S antigens to elicit a protective immune response covering both variants.(37) Clinical trials indicated improved protection compared to non-vaccinated and monovalent-vaccinated groups.(38) Previous studies of nanoparticle-based RBD-displaying vaccines suggested the co-display of mixtures of RBDs on multivalent nanoparticles also elicited broader neutralizing antibodies.(39)

Relatively little investigation has been reported regarding the effect of co-transfection of host cells with more than one mRNA, with most of the focus being devoted to the downstream immune responses. In the case of bivalent mRNA COVID vaccines, where two mRNA sequences may be delivered to the same cell, in addition to displaying both S antigens on the membrane surface, concurrent translation of mRNAs has the potential to result in translated proteins co-assembling into a mosaic S heterotrimer. To our knowledge it has not been determined whether protomers from different SARS-CoV-2 S constructs can co-assemble as a mosaic heterotrimer and whether co-assembly impacts the trimer’s stability, dynamics and antigenicity.

To address these questions from structural and dynamic perspectives, we investigated the nature of spike antigens produced by co-transfection of cells with different combinations of SARS-CoV-2 variants. We demonstrated assembly of combinatorial mixtures of trimers including homotrimers and mosaic trimers assembled by genetically close strains Omicron BA.2 and XBB (OX), as well as distant strains Hu-1 D614G (G614) and Omicron BA.2 (OG) (Figure 1). Hydrogen/deuterium-exchange mass spectrometry (HDX-MS) and biolayer interferometry (BLI) were used to investigate dynamics of the components within the mosaic S assemblies and to determine any impact on RBD structural ordering and receptor binding site accessibility to receptor and antibodies.

**Figure 1.**
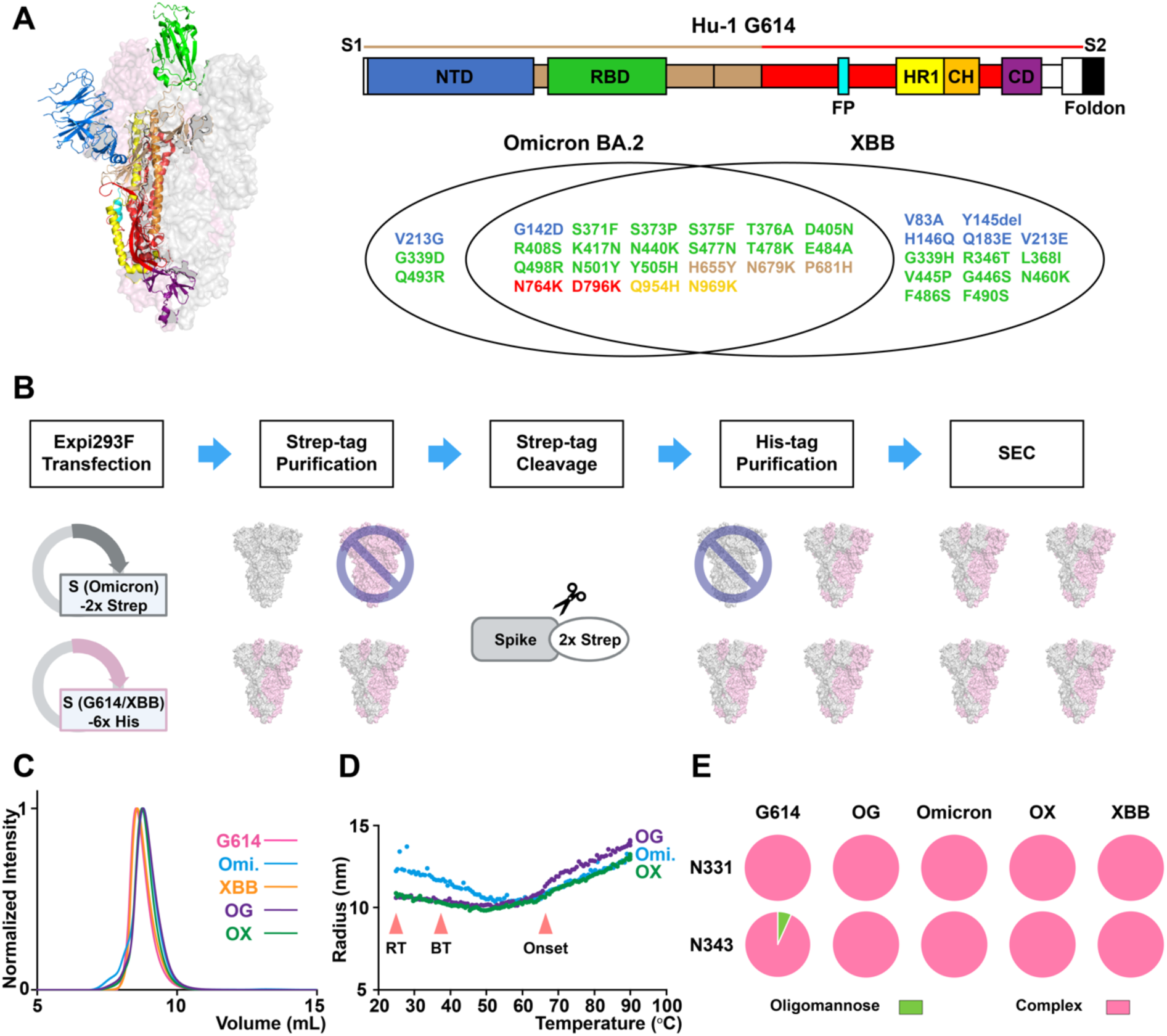
Mosaic S-6P heterotrimer construct sequences, purification process, and characterizations. **(A)** Color-coded S domains on cartoon diagram (PDB: 6VSB) and annotated domain organization. Venn diagram indicates the unique and shared mutations in Omicron BA.2 and XBB constructs with reference to Hu-1 G614 sequence. Color labels match their corresponding domains. S1: receptor-binding subunit; S2: fusion subunit; NTD: N-terminal domain; RBD: receptor-binding domain; FP: fusion peptide; HR1: heptad repeat 1; CH: central helix; CD: connector domain. **(B)** Co-expression and tandem columns purification of mosaic heterotrimer. Strep-tag affinity purification removes homotrimer of construct with only His-tag (G614 and XBB). His-tag affinity purification continues removing the homotrimer of Omicron construct with only Strep-tag. **(C)** SEC chromatograms of homotrimers and mosaic heterotrimers purifications. Major peak eluting at ∼9 mL reveals the trimeric S integrity. **(D)** Thermal stability measurements on Omicron homotrimer and heterotrimers from 25 ^°^C to 90 ^°^C. RT: room temperature, which HDX experiments are performed under; BT: body or physiological temperature; Onset: onset of the unfolding. **(E)** N-glycosylation profiles on all S constructs indicate predominant complex forms at RBD N331 and N343 sites.

Understanding whether such mosaic heterotrimers can form and their implications on the structural properties of S protein is important, as it could influence the way the antigen presents key epitopes to the host immune system, thus impacting vaccine efficacy. These issues are directly relevant to numerous vaccine studies that are pursuing for example, pan-coronavirus or “universal” influenza vaccines that rely upon co-administration of panels of antigen variants in hopes of eliciting protection against highly variable, rapidly evolving viruses.(40, 41) The study we report here provides structural dynamic information to help understand the impact of co-transfection and co-expression on the combinatorial antigens that are produced in these scenarios.

## Results

### Mosaic spike trimers can stably assemble from genetically close and distant SARS-CoV-2 variants

To determine whether co-transfection mimicking bivalent mRNA expression could result in mosaic SARS-CoV-2 S heterotrimer formation, we sought to co-express hexaproline-mutated S ectodomains (S-6P) which were designed to maintain soluble S trimers in the antigenic pre-fusion form (42), from the three selected S variants (Figure 1A). Total S expression levels were compared with equal amounts of total plasmid DNA transfected under nine conditions: Omicron only, G614 only, XBB only, Omicron-G614 co-transfection (2:1, 1:1, and 1:2), and Omicron-XBB co-transfection (2:1, 1:1, and 1:2). Western blot using an antibody targeting a linear epitope in the S2 subunit that is conserved across all three S variants showed that all conditions expressed similar levels of total S protein in the supernatant (Figure S1). This indicates that all three S-6P variants have similar transfection efficiency and expression levels under our experimental conditions, with no significant interference from co-transfection of Omicron and G614 variants, and of Omicron and XBB variants.

To evaluate if mosaic heterotrimers can be formed and how they affect trimer stability, we prepared mosaic spike from Omicron-G614 (OG) and Omicron-XBB (OX) combinations side-by-side with control preparations of Omicron, G614, and XBB homotrimers (Figure 1B, Figure S1). Using the optimized purification protocols and low-passage, viable Expi293F cells, the best yields from parallel purifications of all five constructs were obtained (Table S1). The data agree with the total S expression result and show that the co-expression did not interfere the S trimer production.

Nearly 90% of the protein eluted as a single peak at 8.5-9 mL from the size-exclusion chromatography (SEC) at 4 °C, consistent with well-formed trimers with comparable yields around 4-5 mg (Figure 1C). Both G614 and XBB homotrimers reflect sharp peaks that eluted slightly earlier than Omicron-containing S trimers due to the presence of affinity tags. Omicron homotrimer, eluted later from SEC, showed a modest shoulder eluting slightly before the primary chromatographic peak. The particle hydrodynamic radius and polydispersity measured by dynamic light scattering (DLS) also indicated that Omicron homotrimer had slightly larger radius and polydispersity than other S trimers at 25 °C (Table S2). Thermal melt experiments monitoring particle radius by DLS provided a means of measuring the stability of S assemblies. Both OG and OX mosaic trimers indicated a smaller radius than Omicron homotrimer under room temperature and physiological temperature ranges (Figure 1D). The trimeric integrity and morphology of mosaic heterotrimers were also confirmed by negative-stain electron microscopy (nsEM) images (Figure S1). Therefore, both Omicron and XBB protomers, which possess exactly the same S2 subunit sequence, and Omicron and G614 protomers, which carry numerous mutations around the S1/S2 cleavage site and S2 subunit (Figure 1A), could form mosaic heterotrimers with limited alterations in global trimeric structure and morphology.

Furthermore, based on the yield ratios of the isolated mosaic heterotrimer in the elution to the Omicron homotrimer in the flow-through from the last His-tag purification step (Table S1, OG: 4.7 mg/1.9 mg; OX: 3.6 mg/1.2 mg), the mosaic trimers were estimated to be ∼60% of the total trimer yield from co-transfection, close to the theoretical percentage for stochastic trimer assemblies from 1:1 co-transfection. This suggests that, in both genetically close and distant strains co-transfection, homotrimer formation did not overwhelm heterotrimer formation, and the presence of heterologous S material did not lead to noticeable misassembly or aggregation.

Another important factor that influences immunogenicity of viral glycoproteins is the glycan profile on the displayed antigens. N-linked glycans can contribute to epitopes that antibodies target.(43, 44) Additionally, in SARS-CoV-2 S, N-linked glycans on the RBD have been suggested to affect trimer open/closed state equilibria, which could modulate exposure of some epitopes.(45) To determine how mosaic S heterotrimers glycoprofiles compare with their respective homotrimers, we carried out glycan profiling for the set of mosaic and homotrimer spikes. Our results show that both N-glycosylation sites on the RBD are predominantly complex forms across all five trimeric assemblies (Figure 1E). The glycan profiles we determined in our constructs are consistent across purification batches and in agreement with previous studies on the soluble pre-fusion S ectodomain constructs (46) as well as protein nanoparticle vaccines featuring pre-fusion S or RBD antigens (31, 47) that have been shown to elicit potent neutralizing antibody responses. For the rest of the N-glycosylation sites we characterized, the glycan profiles are mostly unaltered in mosaic S trimers except for the N801, which exhibited more processing towards complex forms in OG heterotrimer (Figure S2).

### Dynamic features of Omicron and XBB protomers in the mosaic heterotrimer are similar to their homotrimer counterparts

Next, we carried out HDX-MS structural dynamic analysis comparing three S-6P homotrimers: G614, Omicron and XBB in order to obtain baseline structural dynamic profiles against which we could compare structural dynamics and trimer integrity profiles of the OX and OG mosaic heterotrimers.

HDX-MS analysis on G614, Omicron and XBB homotrimers revealed isolate-specific differences in conformational preferences and S2 dynamics highlighting regions of the polypeptide that were ordered differently among these three homotrimer antigens (Figure 2A, Figure 2B, Figure S3). In our previously reported comparison of S-2P trimers (13), peptides 388-392 and 982-990 were found to serve as effective reporters for RBD up/down configurations as increased dynamics near the C-terminus of the RBD facing the apex of central helices (peptide 388-392) and the central helical apex (peptide 982-990) were observed when S adopts an open conformation. In the present experiments, both of these peptides indicated that dynamics were greater in G614 than in Omicron, which was also significantly greater than in XBB (Figure 2C, peptide #1 and #2). When S is in its open state, the RBD-up conformation also affects dynamics in the S1 hinge region connecting RBD to the rest of the spike trimer. For example, the N-terminal peptide 516-533 connecting to the RBD, showed increased dynamics in response to RBD shifting up, while peptide 542-568, was more protected by the neighboring protomer facing towards the trimer core (Figure 2C, peptide #3 and #4). These dynamic changes all support the conclusion that the G614 S-6P trimer exhibits a greater conformational bias toward populating an open state than Omicron and XBB S-6P trimers.

**Figure 2.**
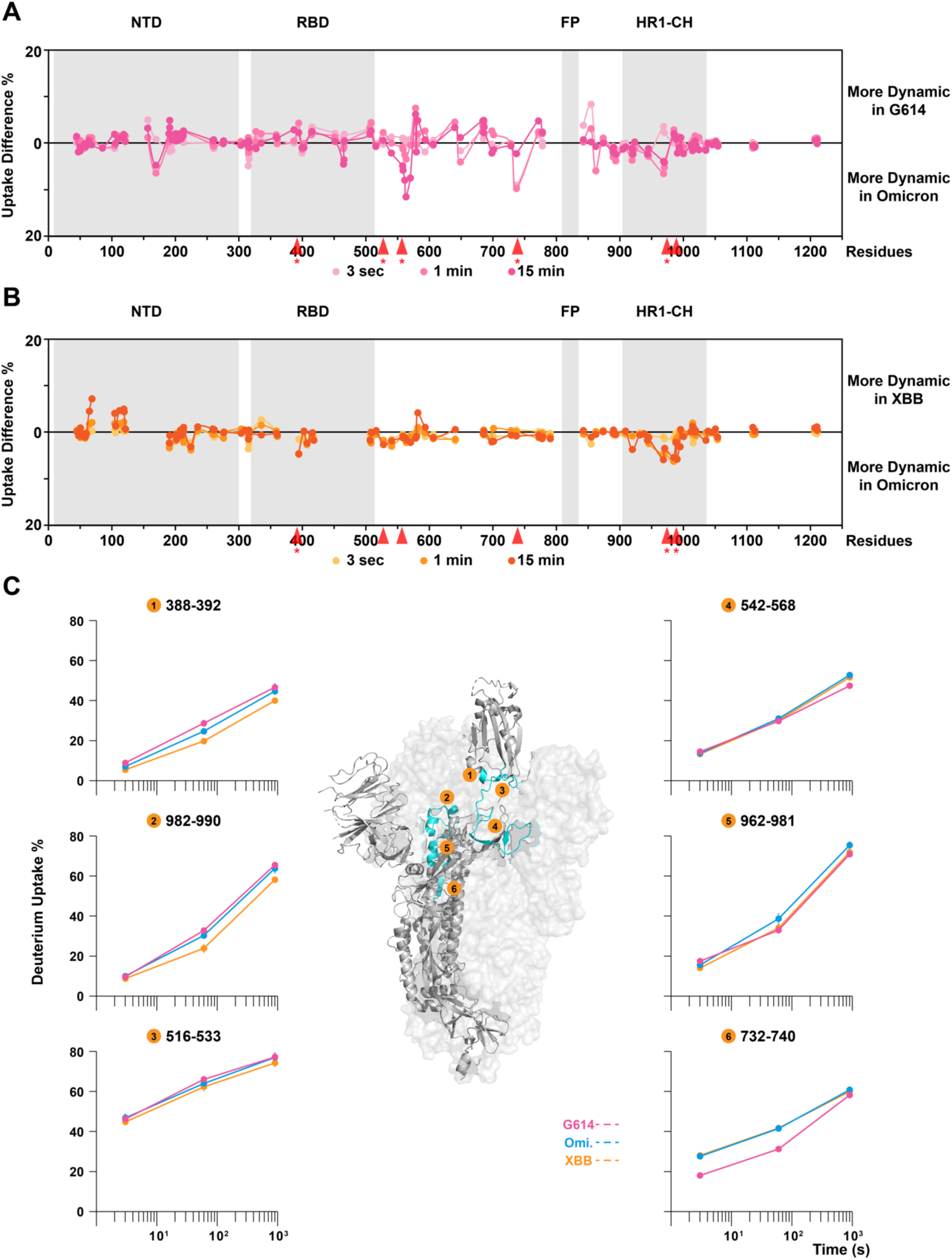
Structural dynamic comparisons between G614, Omicron, and XBB S-6P homotrimers. Differential deuterium uptake plots of **(A)** G614 versus Omicron and **(B)** XBB versus Omicron with **(C)** zoom-in homologous peptide comparisons in the region exhibiting major differences, reveal the strain-specific dynamic profiles. Peptide locations are indicated by red arrows in panel A and B. Asterisk next to the red arrow indicates statistically significant difference between uptake levels on that peptide. Error bars (SD from n > 3) less than 1.5% were masked by the data points. PDB: 6VSB.

Sequence variations in the N-terminal domain (NTD) and RBD, especially in the receptor-binding motif (RBM) regions among the three constructs resulted in different digestion patterns, limiting the set of homologous peptides that could provide direct comparisons in these two regions (Figure 2B). By contrast, Omicron and XBB share the same S2 sequences and digestion patterns. HDX-MS analysis revealed that the only significant dynamic differences in the S2 subunit were observed in the heptad repeat 1 (HR1) region reflecting a less open state in XBB (Figure 2C, peptide #2 and #5, Table S3). Residue substitutions around the S2’ and HR1 region in Omicron and XBB relative to the ancestral Hu-1 G614 variant yielded differences in S2 subunit dynamics near the S2’ site, where both Omicron and XBB showed increased deuterium uptake over G614 (Figure 2C, peptide #6, Table S3). Overall, summarizing the isolate-specific dynamics in G614, Omicron and XBB S homotrimers: G614 S adopted more open conformations with stabilized S2 subunit; Omicron S showed less open conformations than G614 but possessed greater dynamics in the S2 subunit from the HR1 domain extending to the fusion peptide-proximal region (FPPR); and XBB S indicated the least open state preferences and compromised S2 subunit dynamics. The dynamic features in G614 and Omicron S closely agree with previously reported results.(48)

To understand whether the formation of mosaic heterotrimers could alter the conformational preference, structural dynamics, and epitope exposure of these antigens, we compared the HDX-MS results from homologous peptides in the mosaic S heterotrimer with those in the corresponding homotrimers. Since the homologous peptides with identical sequence from two constructs will co-elute in HDX-MS, a multi-modal mass envelope is anticipated if a given peptide behaves differently in the different protomers. However, in the cases when the separation was not great enough to deconvolute the bimodal populations into individual protomer contributions, a broadened mass envelope reflecting the average of two underlying populations is observed.

In the Omicron-XBB (OX) mosaic S trimer, the Omicron and XBB protomers mostly retained the dynamic features of their respective homotrimers (Figure 3, Figure S4). The dynamics of NTD and RBD from each protomer were not significantly affected by mosaic trimer formation as can be seen in the differential plot compared against the baseline Omicron homotrimer (Figure 3A). The peptide uptake profiles in the NTD of OX reflect the average of corresponding uptake levels in Omicron and XBB homotrimers (Figure 3B, peptide #1 and #2).

**Figure 3.**
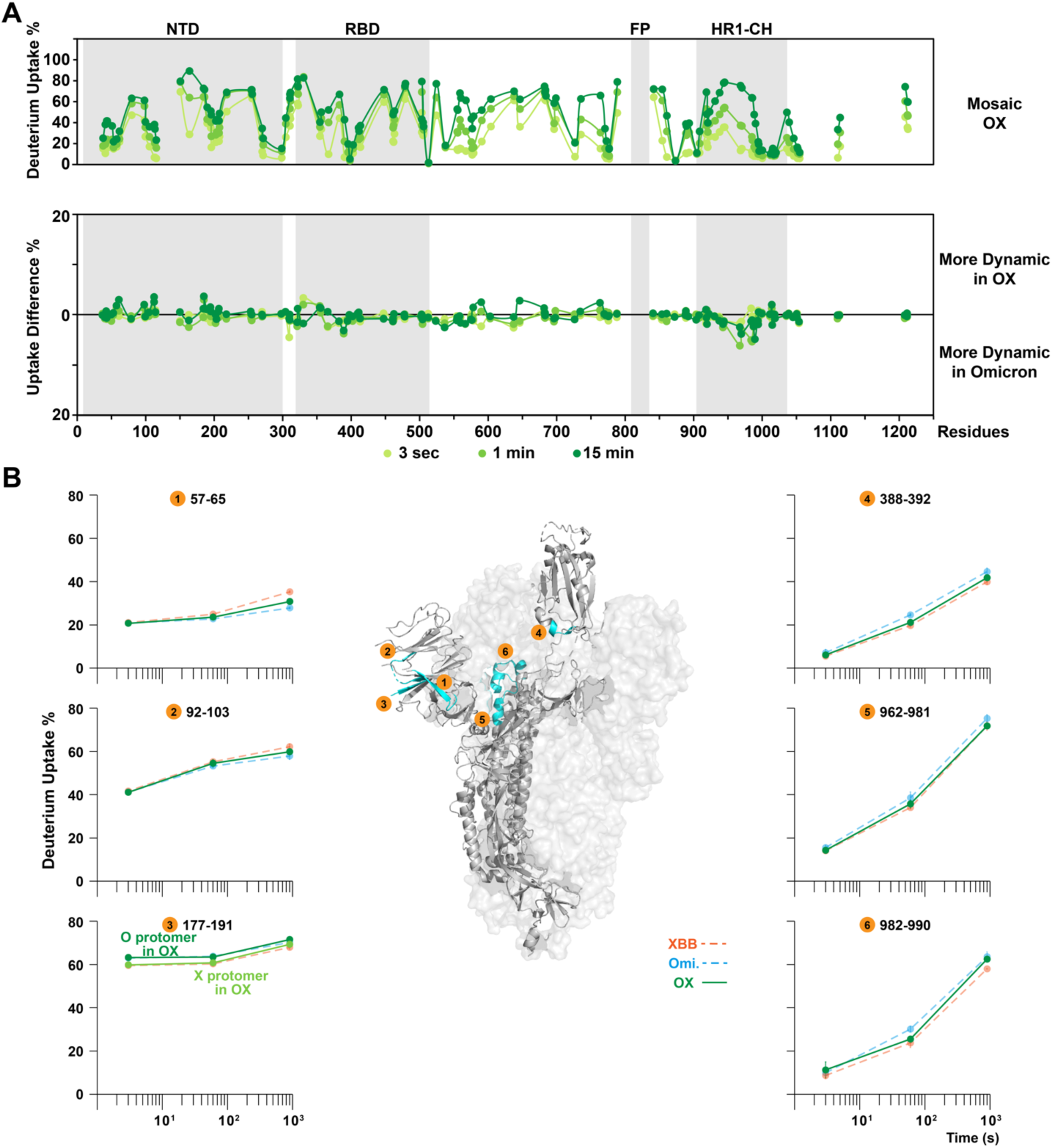
Structural dynamic differences upon forming mosaic Omicron-XBB heterotrimers. **(A)** Deuterium uptake plots and differential plots of Omicron-XBB mosaic heterotrimer and **(B)** pair-wise comparisons in homologous peptides, depict dynamic features of mosaic OX heterotrimers. Peptide #1-3 featuring NTD, reveal the NTD dynamics are retained as in homotrimers. In peptide #3, the darker green plot indicates the behavior of Omicron protomer in the OX mosaic trimer while the lighter green plot indicates the behavior of XBB protomer in the OX mosaic trimer. Peptide #4-6 showed the dynamic impacts on spike conformational states, where OX has the intermediate uptake levels. Error bars (SD from n > 3) less than 1.5% were masked by the data points. PDB: 6VSB.

Homologous peptides such as the peptide spanning residues 177-191 (Figure 3B, peptide #3), where the Q183E substitution in the XBB sequence produced peptides that differ by the single amino acid substitution, could also be distinguished in the mass spectrometer and directly compared with counterparts in the homotrimers. The peptide in the Omicron protomer of the OX mosaic trimer showed the identical dynamic profile as in the Omicron homotrimer and the peptide in XBB protomer matched the XBB homotrimer peptide as well (Figure 3B, peptide #3). These examples thus demonstrated that Omicron and XBB protomers mostly retained the isolate-specific dynamic features over the immunodominant NTD and RBD domains.

Besides the local structural orderings, overall, the dynamic profile of the mosaic OX trimer was found to present as a hybrid between Omicron and XBB (Figure 3). Regions that are structurally associated with S open/closed states such as the RBD C-terminus (Figure 3B, peptide #4) and apex of the central helix (Figure 3B, peptide #5 and #6) exhibited dynamics intermediate between the two homotrimers. The conservation of isolate-specific dynamics in the NTD and RBD as well as conformational preferences indicates relatively little crosstalk and alteration in behavior of the heterologous protomer in the OX co-assembled spikes, most likely due to the commonality of their S2 subunits.

### A greater degree of crosstalk is observed between protomers in the Omicron-G614 mosaic heterotrimers

We next investigated the dynamic behavior in the Omicron-G614 (OG) mosaic heterotrimer. This combination enabled us to examine whether a mosaic S trimer with divergent S2 subunits may exhibit more prominent effects on inter-protomer packing or allosteric effects across the trimeric spike assembly that could affect their antigenicity. In the OG mosaic trimer, similar effects were observed in the RBD as in the OX case, which exhibited uptake levels intermediate between those in Omicron and G614 homotrimers (Figure 4B, peptide #1 and #2, Figure S5). Peptide 471-486, spanning the loop connecting the RBD core and the RBM located at the surface of RBD, is a target for neutralizing antibodies and is highly mutated during viral evolution, conferring immune escape.(49, 50) The S477N, T478K, and E484A substitutions in peptide 471-486 observed in Omicron vs G614 protomers enable the homologous peptides to be distinguished, allowing direct comparison between the behavior of this peptide in the two types of protomers. In this case, the uptake level in each protomer matched the behavior seen in the respective homotrimer (Figure 4B, peptide #3). Notably, in contrast to the conformational preferences observed in the OX heterotrimer, the OG heterotrimer appeared to favor open conformations. Specifically, the peptides reporting on RBD configurations in the OG heterotrimer more closely paralleled their behavior in the G614 homotrimer, showing the dominant influence of the G614 protomer on the mosaic OG assembly (Figure 4B, peptide #1, #2 and #6, Figure S5). One of the exceptions to this G614 dominance was observed in the hinge region (Figure 4B, peptide #4), where the dynamic behavior of this peptide in OG heterotrimer more closely recapitulated its behavior in the Omicron homotrimer. While increased dynamics at C-terminal RBD (peptide #2) and helical apex (peptide #6) were consistent with greater exposure in an open conformation, the dynamics of the peptide 553-568 loop, which is located at the interface between RBD and a neighboring NTD, was found to be more affected by the positioning and packing of the NTD on the neighboring protomer (Figure 4B, peptide #4). The dynamic features at this site in the OG heterotrimer mirrored the behavior in the Omicron homotrimer indicating that the NTDs maintained a similar position and conformation as in Omicron homotrimer, despite the RBD favoring the more open state in the same OG mosaic trimer. In the mosaic trimer, these structural motifs appear to be somewhat decoupled in their dynamic behavior.

**Figure 4.**
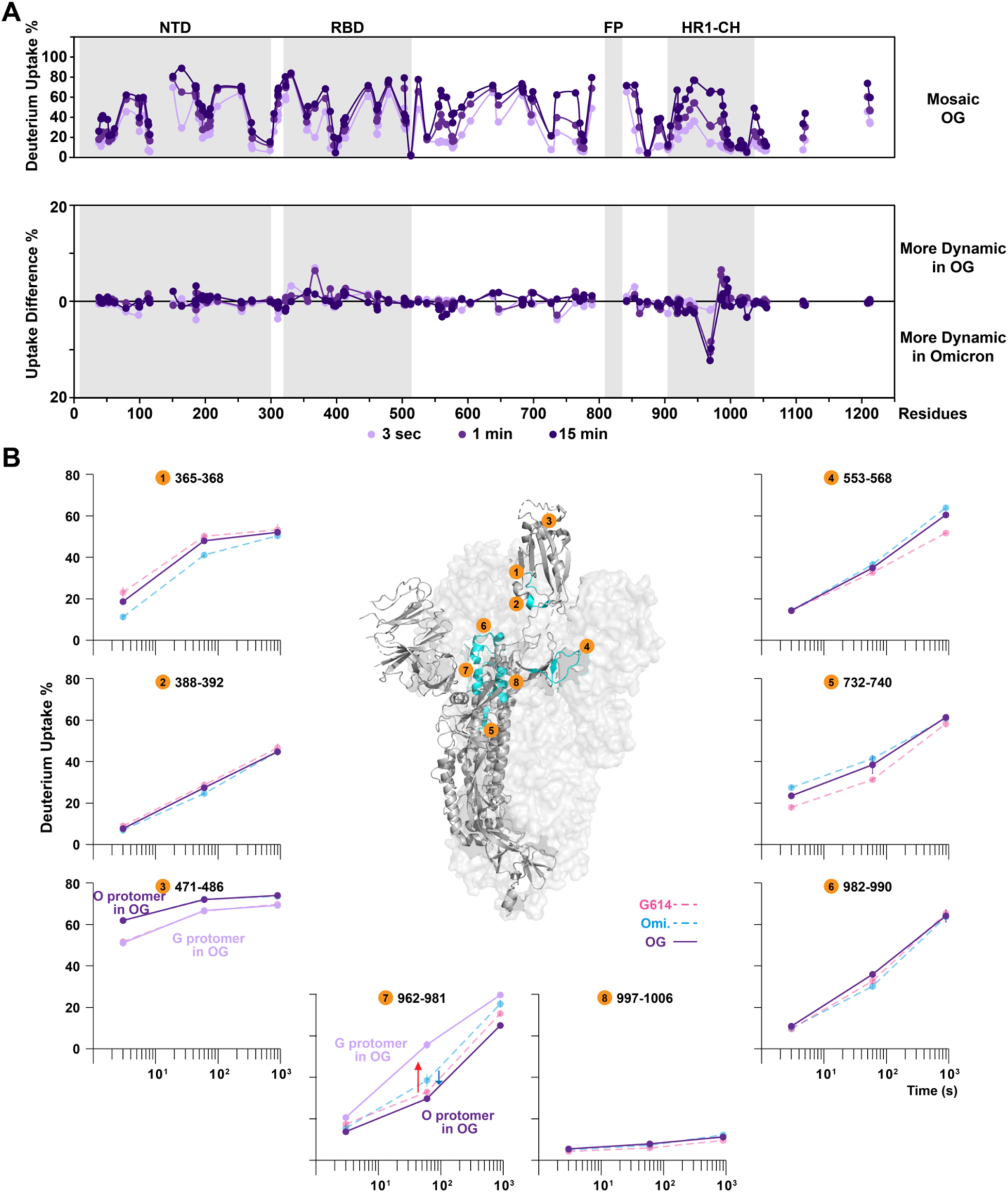
Dynamic differences upon forming mosaic Omicron-G614 heterotrimers. **(A)** Deuterium uptake plots and differential plots of Omicron-G614 mosaic heterotrimer and **(B)** pair-wise comparisons in homologous peptides, depict dynamic features of mosaic OG heterotrimers. Peptide #1-3 featuring RBD, reveal the RBD dynamics are retained as in homotrimer. Peptide #4 is on the S1 hinge region, interacting with neighboring NTD, shows similar dynamic profile to Omicron. Peptide #5-7 indicate the increasing dynamics over FPPR in the OG heterotrimer. In peptide #3 and #7, the darker purple plot indicates the behavior of Omicron protomer in the OG mosaic trimer while the lighter purple plot indicates the behavior of G614 protomer in the OG mosaic trimer. Peptide #8 confirms the spike integrity reflected by the well-protected central helix core. Error bars (SD from n > 3) less than 1.5% were masked by the data points. PDB: 6VSB.

Prominent dynamic changes in the OG heterotrimer were found in the FPPR region between S1/S2 and S2’ cleavage sites (Figure 4B, peptide #5) and HR1 domain (Figure 4B, peptide #7). Notably, homologous peptides in the region 962-981 were differentiated from two protomers due to the N969K mutation in Omicron. This helix-loop-helix peptide was intrinsically more dynamic in the Omicron homotrimer than the G614 homotrimer, in agreement with a previous study (48), but the trend was reversed in the Omicron protomer and the G614 protomer when they co-assembled (Figure 4B, peptide #7). Examining the mass spectra revealed a bimodal mass envelope for this FPPR peptide in the Omicron homotrimer at the 1-minute time point (Figure 5) with a fast-uptake population in the Omicron homotrimer that could adopt a more exposed, dynamic conformation. However, the same region in the Omicron protomer in the mosaic OG trimer showed decreased uptake levels with less detectable peak broadening and suppressed bimodal spectra which mostly resembled the slow-uptake population in the Omicron homotrimer (Figure 5). This suggests that the FPPR in the Omicron protomer of the OG mosaic trimer stably maintains its ordered conformation. On the other hand, the homologous peptide in the G614 protomer showed the opposite trend, exhibiting significantly increased dynamics, in the OG mosaic trimer (Figure 5). Enhanced dynamics in the G614 protomer of the OG mosaic trimer was reflected by the presence of novel dynamic conformations in FPPR, that gave rise to a bimodal mass envelope at the 1-minute time point compared to the unimodal ordered conformation in G614 homotrimer (Figure 5). This phenomenon indicates that Omicron-specific mutations Q954H and N969K can allosterically induce the neighboring protomer to adopt highly dynamic conformations in the FPPR.

**Figure 5.**
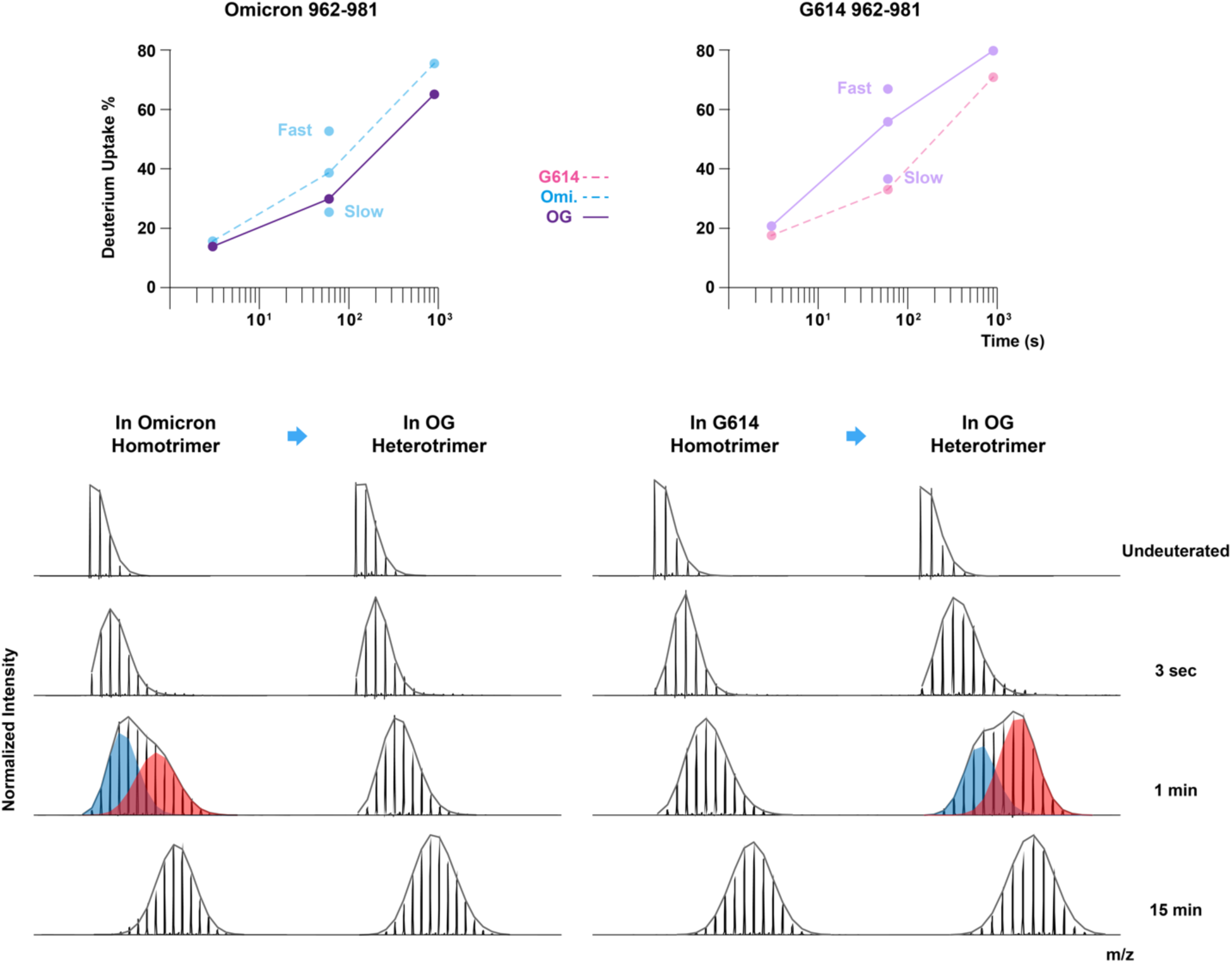
Protomer-specific dynamics near S2 subunit apex in the Omicron-G614 heterotrimers. Peptides 962-981 containing a N969K mutation in Omicron construct is used to differentiate deuterium uptake profiles between Omicron and G614 protomers. Dynamic changes in Omicron protomers (left panel) and G614 protomers (right panel) from their homotrimers to OG heterotrimer are compared. Fast and slow uptake levels labeled on the plots, correspond to red (fast) and blue (slow) populations of the bimodal mass envelopes at 1-minute time point.

The crosstalk between protomers in the OG mosaic heterotrimer gave rise to novel structural and dynamic features that S homotrimers did not present. Specifically, the mosaic heterotrimer favored an open state as seen in the G614 homotrimer, with the isolate-specific orderings in NTDs and RBDs for the Omicron protomers mostly retained. The pre-fusion S helix-loop-helix region at FPPR became more ordered in the Omicron protomers with the induction of highly dynamic conformation in the G614 protomer over the same region. Despite such prominent dynamic effects in the S2 subunit, the OG mosaic trimer integrity and stability were well-maintained, as indicated by constantly protected central helical domains from HDX-MS (Figure 4B, peptide #8, Figure S5), thermal stability results (Figure 1D), and nsEM images (Figure S1). Thus, the sequence differences in S2 between the ancestral G614 subvariant and prevalent Omicron subvariant were not sufficient to impact the trimer’s ability to assemble but did result in modulation of the trimer’s dynamic profile at positions across the spike assembly.

### Antigenic effects of mosaic trimer formation

Previous studies have reported that over 90% of neutralizing antibodies targeted RBD and NTD.(14–16) Retaining the structural ordering and functioning in NTD and RBD is important for assessing the antigenicity as well as vaccine efficacy. To further investigate whether the localized structural dynamic differences in the mosaic trimers have antigenic effects, bio-layer interferometry (BLI) was used to compare human ACE2 (hACE2) and antibody binding kinetics for mosaic versus homotrimer spikes.

In the dimeric hACE2 binding experiments, all five constructs showed strong binding affinities with K_D_<10 nM (Figure 6), agree with previously reported K_D_ measured by surface plasmon resonance.(51) We then tested binding with monoclonal antibody (mAb) C68.13 that competes with ACE2 binding and has been reported to exhibit 2-3 orders of magnitude weaker inhibition of XBB that G614 and Omicron.(52) This allows us to selectively examine how the critical RBD epitope presentation is modulated on the mosaic heterotrimer. In our binding assay, as expected, the G614 homotrimer, Omicron homotrimer and OG mosaic trimer were found to bind C68.13 with high affinity, K_D_ ∼1 nM (Figure 6). Flat dissociation curves resulted in k_off_ rates in lower 10E-4 levels, close to the detection limit, indicating very stable binding. The XBB homotrimer bound C68.13 with K_D_ ∼95 nM, 50-to-100-fold weaker binding to this mAb than G614 and Omicron homotrimers. The mosaic OX showed comparably strong binding from almost 5-fold k_off_ rate changes from XBB homotrimers (Figure 6). This similar k_off_ rate to the Omicron and G614 homotrimers’ suggested the mode of interaction between the RBD and antibodies were relatively unaltered by the presence of adjacent protomers with weak binding affinities.

**Figure 6.**
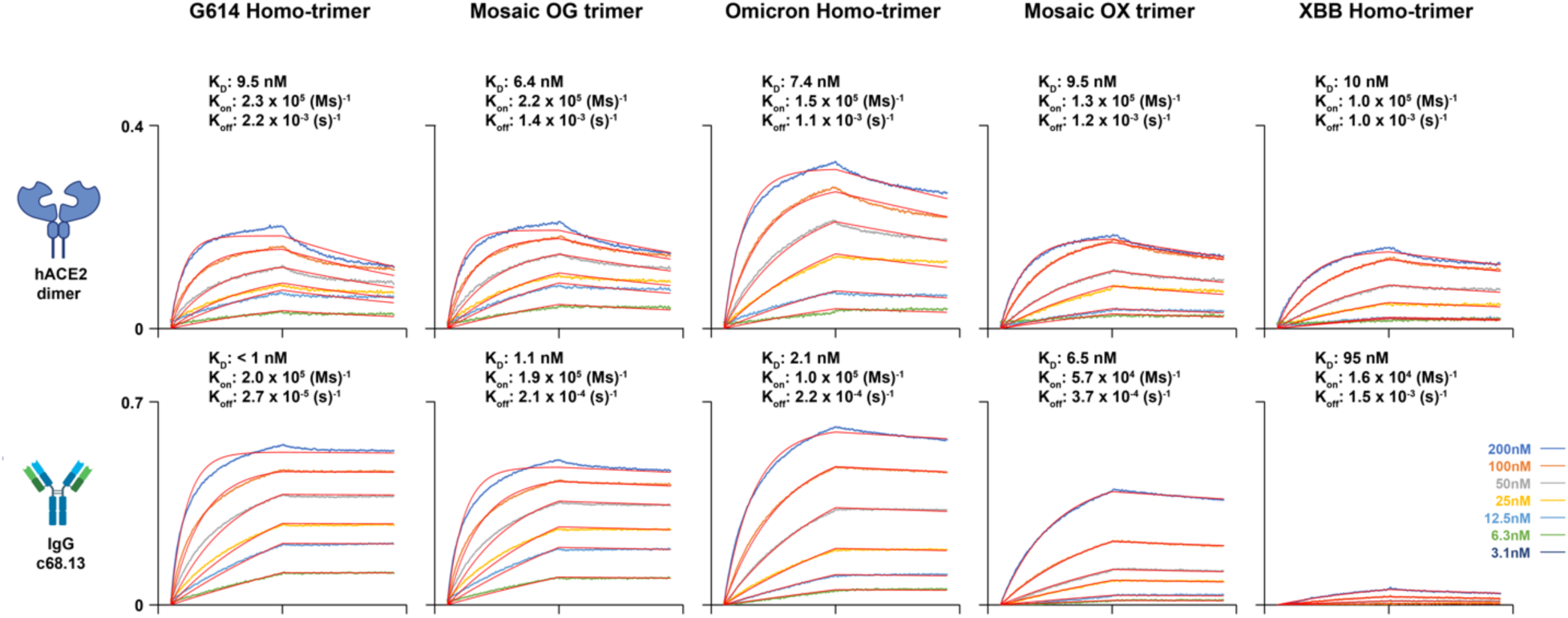
RBD structural orders of mosaic S-6P heterotrimers. BLI measures K_D_ on five spike trimers interacting with dimeric human ACE2 and monoclonal antibody c68.13. Binding curves are aligned at the start of association step and fitted globally with a 1:1 ligand binding model.

In both hACE2- and selected mAb-binding cases, the structural orderings at RBM interface of mosaic S heterotrimer were shown to be well-conserved. Our BLI results align with the inferences from the HDX experiments, suggesting that mosaic heterotrimers (OX and OG) impose limited restriction on RBD dynamics and receptor/antibodies binding.

## Discussion

Glycoproteins on enveloped viruses such as influenza, HIV, and coronaviruses are the primary antigenic features on the virus surfaces. The stability, dynamics, and structural ordering of these multimeric assemblies influence the display of antibody epitopes and their recognition by neutralizing antibodies. Viruses with RNA genomes including SARS-CoV-2 evolve rapidly when spreading through large populations and adapting to escape host immune responses. In order to combat emergence of such viral diversity and maintain or increase breadth of immune responses, many strategies under investigation combine antigens from multiple variants and viral strains into polyvalent antigen cocktails. In the case of nucleic acid-launched vaccines such as those employing mRNA, immunization with mixtures of genes encoding the antigens from various strains are delivered to the vaccinee such that co-transfection of cells has the potential to lead to co-expression of two or more antigen variants. As we have shown here, this also can lead to co-assembly of mosaic antigens. Such a strategy of combining a bivalent antigen cocktail was used for a COVID-19 booster immunization, deployed in 2022, which combined proline-stabilized, pre-fusion Hu-1 and Omicron variants.(37) Of note, such a situation of co-expression and co-assembly of antigen variants may also take place if a cell is co-infected with more than one strain of virus during superinfection of the host.

The three SARS-CoV-2 S variants examined here: Hu-1 G614, Omicron BA.2, and XBB, were selected based on their genetic distances and significance as vaccine immunogens from distinct waves of the pandemic. The G614 variant, close to the ancestral Wuhan-Hu-1 WT and first detected in early 2020, led to subsequent infection waves worldwide before major vaccination was deployed.(6) In contrast, Omicron subvariants and XBB emerged in late 2021 and 2022, respectively, carry mutations that emerged after the administration of two or three vaccine doses in most developed countries and widespread natural infection worldwide.(19, 21) The genetic distance between G614 and Omicron is relatively large compared to the distance between Omicron and XBB (Figure 1A). In particular, XBB is a recombinant variant formed from two Omicron subvariants, resulting in notably different S1 but identical S2 subunit sequences compared to Omicron BA.2. Our choice of S variants thus enabled us to investigate the impacts of the bivalent mRNA vaccine-expressed antigens as well as the structural and dynamic impact of S2 subunit sequence similarity and differences on mosaic antigen display.

We found that mosaic S trimers from co-transfection maintained the overall protein integrity and morphology relative to the corresponding homotrimers. The yield of mosaic trimer closely matched the theoretical yield for respective homotrimers, suggesting a negligible cost for co-expressing and co-assembling protomers from different variants into mosaic trimers. We speculate that this well-ordered S morphology is due to the highly similar S2 subunit sequence and conformation. We also note that the hexaproline-stabilized form of the S ectodomain may suppress variant-specific effects impacting trimer co-assembly, such as the different intrinsic propensities to convert to a post-fusion form and conformational switching.(53, 54) The incorporation of proline mutations improved overall purification yields and experimental consistency. We anticipate that the observed effects we describe may be even more prominent for less stabilized diproline, S-2P, trimers that are used in the approved COVID vaccines.

The mosaic heterotrimers we expressed should exist as a mixture of AAB and ABB heterotrimer forms (A and B represent two different protomers) resulting from the 1:1 ratio of transfected constructs. Western blots showed a roughly 1:1 ratio of each heterotrimer form (Figure S1). Therefore, the deuterium uptake level from HDX-MS reflected a compounded dynamic profile of both AAB and ABB forms. Our conclusion of mosaic heterotrimer dynamics thus likely underestimates specific effects as we could not differentiate between protomer behavior in AAB- or ABB-specific individually.

Numerous on-going investigations seek to generate immune responses that can provide broad protection against rapidly evolving viruses. Such efforts on pan-coronavirus or universal influenza vaccine studies seek to elicit broad responses by vaccinating with cocktails of antigen variants in some cases co-formulated and in others co-displayed on multivalent nanoparticle platforms.(39, 55–58) One hope of such polyvalent display of antigenic variants is to select for antibodies that are able to recognize the conserved features while tolerating variability in other regions through avid binding of multivalent nanoparticles to B-cell receptors. In the case of the mosaic trimers we have analyzed, it may similarly be possible for hetero-protomers in a given co-assembled trimer to engage with B cell receptors that can bivalently bind to conserved epitope features displayed in two distinct contexts, helping to favor a broad response.

Our experiments have shown that, at least in the SARS-CoV-2 spike system, bivalent co-formulation and transfection of S variants leads to the formation of mosaic heterotrimers that retain structural integrity while exhibiting novel dynamic behaviors distinct from the parental homotrimers. Our experiments demonstrated interprotomer crosstalk especially in mosaic spike heterotrimers from relatively divergent sequences. For future vaccine designs it may be necessary to take into account the potential formation of mosaic heterotrimers and resulting structural and immunological implications following polyvalent nucleic acid vaccination such as for pan-coronavirus and universal influenza vaccines or in targeting highly variable viruses such as HIV.

## Acknowledgements

We thank Jamie Guenthoer (Fred Hutchinson Cancer Center) for providing c68.13 antibody as a ligand in BLI experiments to test binding kinetics. We also thank Jacob Croft (U. Washington) for assistance in setting up nsEM image collection. The proteomics work was supported by Mass Spectrometry Center in the School of Pharmacy (U. Washington). This work was supported by NIH grants R01-AI156808 to K.K.L and R01-AI38709 to J.O. and by S10-OD030237.

## Data Availability

Data are available in the manuscript and supporting information files. Both the unprocessed HDX-MS and N-glycosylation analysis in raw data format and the processed excel file showing peptide identification and relative uptake levels have been deposited to the ProteomeXchange Consortium via PRIDE partner repository with accession number PXD060785. (FOR REVIEW ONLY: please use the token WxlBOSMRlUce to access the data)

## Materials and Methods

### Plasmid construction

The gene sequences encoding SARS-CoV-2 S ectodomain with hexaproline mutations (S-6P: F817P, A892P, A899P, A942P, K986P and V987P)(42), GSAS substitution at S1/S2 site (RRAR) and C-terminal T4 fibritin foldon were cloned into pCMV plasmid. The open reading frame (ORF) sequence of Hu-1 G614 S-6P construct is the reference Wuhan-Hu-1 sequence (residues 14-1211) with the D614G mutation, C-terminal His-tag and the alterations mentioned above. Omicron BA.2 construct was assembled from the ORF in the pαH Omicron BA.2 S-2P plasmid (Addgene plasmid #184829) into pCMV plasmid and mutated with four additional prolines using HiFi DNA Assembly (NEB). XBB construct was assembled from the ORF in the pVRC8400 XBB S-2P plasmid (Addgene plasmid #160474) into pCMV plasmid and mutated with four additional prolines and a C-terminal His-tag (NEB).

### Transient transfection

293T adherent cell line was used for optimizing co-transfection condition. Cells were split into 12-well plate (1 mL per well) at seeding density (0.1 × 10^6^ cells per well). Confluent cells (0.5 × 10^6^ cells per well) were transfected with total 2 μg DNA and 2 μL Lipofectamine™ 3000 (Invitrogen) based on Lipofectamine™ 3000 reagent protocol. DNA being transfected or co-transfected into the cell cultures under nine conditions: Omicron only, G614 only, XBB only, Omicron-G614 co-transfection (2:1, 1:1, and 1:2), and Omicron-XBB co-transfection (2:1, 1:1, and 1:2). After 2 days in the 37 °C incubator (5% CO_2_), 200 μL supernatant from each well was collected for SDS-PAGE to check expression level.

Suspension Expi293F cell culture was used for spike protein expression. For a 700 mL cell culture at confluency (3 × 10^6^ cells/mL), 0.7 mg total DNA (1:1 for co-transfection) and 2.1 mL ExpiFectamine™ 293 reagent were mixed in Gibco Opti-MEM and added into the cell culture following ExpiFectamine™ 293 user guide. The transfected culture was incubated at 37 °C and 5% CO_2_ with constantly shaking at 125 RPM for 5 days before harvest.

### Protein purification

On the fifth day post-transfection, Expi293F cells were harvested by centrifugation at 2000 RCF for 30 minutes. Supernatant was vacuum-filtered through 0.45 μm αPES filters and supplemented with Tris-HCl (pH 8.0) and NaCl.

For G614 and XBB spike homotrimers purification, imidazole was added to the supernatant to reach a final concentration of 50 mM Tris-HCl (pH 8.0), 200 mM NaCl and 20 mM imidazole, same as the Ni-IMAC binding buffer composition. A 2 mL Ni-IMAC resin (Thermo Fisher Scientific) was washed and equilibrated with binding buffer and packed for a gravity flow column. Supernatant was loaded and flowed through this gravity flow column, followed by four times 10 mL (5 CV) wash with binding buffer. The bound protein was then eluted from the column with three times 6 mL (3 CV) Ni-IMAC elution buffer (50 mM Tris-HCl, pH 8.0, 200 mM NaCl, 300 mM imidazole). Elution fractions were SDS-PAGE checked, and then combined. Combined spike protein solution was buffer-exchanged to reduce the imidazole level below 1 mM and concentrated to ∼1 mg/mL with an Amicon Ultra-15-mL-centrifugal filter (30 K, Millipore), followed by flash frozen in liquid nitrogen and stored in -80 °C freezer.

For Omicron, mosaic Omicron-G614, and mosaic Omicron-XBB trimers purification (Figure 1B), ethylenediaminetetraacetic acid (EDTA) was added to a reach a final concentration of 100 mM Tris-HCl (pH 8.0), 150 mM NaCl and 1 mM EDTA, same as the Strep-Tactin^®^ XT wash buffer composition. A 3.5 mL Strep-Tactin^®^ XT 4Flow^®^ high capacity resin was washed and equilibrated with wash buffer and packed for a gravity flow column. Supernatant was loaded and flowed through this gravity flow column, followed by four times 8.5 mL (2.5 CV) wash. The bound protein was then eluted from the column with three times 5 mL (1.5 CV) Strep-Tactin^®^ XT elution buffer (100 mM Tris-HCl, pH 8.0, 150 mM NaCl, 1 mM EDTA, 50 mM biotin). Elution fractions were combined, buffer-exchanged to reduce the biotin level below 1 mM and concentrated to a final volume below 1 mL. HRV 3C protease (Pierce™) was added accordingly based on the protein solution volume for the digestion to remove TwinStrep tag overnight at 37 °C. On the next day, digested samples were spun at 15000 RCF for 10 minutes to remove possible protein aggregation pellet. The Omicron homotrimer protein could now be flash frozen in liquid nitrogen and stored in -80 °C freezer. For both mosaic heterotrimer purifications, samples were flowed through a 1 mL pre-equilibrated prepacked StrepTrap™ XT column (Cytiva) at 1 mL/min to separate undigested protein. Flow-through was reloaded to the column for better purification results. The final flow-through containing heterotrimer with TwinStrep tag removed was collected and buffer-exchanged to the nickel column binding buffer. Samples were flowed through a 1 mL pre-equilibrated prepacked HisTrap™ HP column (Cytiva) at 1 mL/min, followed by 7 mL (7 CV) wash. The bound protein was then eluted from the column with 3 mL (3 CV) elution buffer. SDS-PAGE checked elution fractions were combined, buffer-exchanged to reduce the imidazole level below 1 mM and concentrated to ∼1 mg/mL with an Amicon Ultra-15-mL-centrifugal filter (30 K, Millipore). The duo-affinity-column purified mosaic spike heterotrimers were then flash frozen in liquid nitrogen and stored in -80 °C freezer.

All the spike trimers were finally size-exclusion chromatography (SEC) purified before major experiments. 500 μL sample was thawed and spun at 15000 RCF for 10 minutes to remove possible protein aggregation pellet before injecting to Superdex 200 column (GE) on an AKTA Pure system (GE). With the SEC running buffer (10 mM HEPES, 200 mM NaCl, 0.02% NaN_3_, pH 7.5) and flow rate at 0.5 mL/min, spike trimer was eluted at around 8.5-9 mL. The SEC purified spike trimers were concentrated to desired concentrations according to the experiment need.

### Dynamic light scattering

Dynamic light scattering measurements were performed on a DynaPro NanoStar II (Wyatt) to characterize spike trimer integrity, homogeneity and thermal stability. SEC purified spike samples at 0.5 mg/mL were first spun down at 15000 RCF for 20 minutes to remove aggregation pellet. 5 μL of the sample was then injected into a 2-μL quartz cuvette (Wyatt). Each DLS run was measured with twenty 10-second acquisitions at 25 °C by LASER auto-attenuation. Thermal melt measurements were performed from 25 °C to 90 °C with the rate at 0.5 °C /min. Five measurements were taken and averaged at each temperature point.

### Hydrogen/Deuterium-exchange mass spectrometry

For the hydrogen/deuterium-exchange (HDX) experiments, 10 μg each of Hu-1 G614, Omicron BA.2, XBB, mosaic Omicron-G614 and mosaic Omicron-XBB spike trimer was incubated in the deuteration buffer (10 mM HEPES, pH* 7.5, 85% D_2_O, Cambridge Isotope Laboratories, Inc.) at 22 °C for three different timepoints: 3, 60, 900 seconds (five replicates for 3 seconds and four replicates for 60 and 900 seconds). All exchanged samples were immediately mixed with an equal volume of ice-chilled quench buffer containing 8 M urea, 200 mM tris(2-chloroethyl) phosphate (TCEP) and 0.2% formic acid (FA) to pH 2.5, transferred into an autosampler vial with metal cap and flash frozen in liquid ethanol. Samples were loaded to the LC-MS system by an automating platform and analyzed by a Synapt G2 mass spectrometer (Waters) with the settings described in previous publication.(59) Samples were in-line digested into peptides by an immobilized pepsin column (2.1 × 50 mm) and loaded onto CSH C18 trap cartridge (Waters) with loading buffer (2% acetonitrile (ACN), 0.1% trifluoroacetic acid (TFA)) at 800 μL/min. Peptides were then separated by a UPLC CSH C18 column (1.0 × 50 mm, 1.7 μL, Waters) with a 15-min linear gradient from 3% to 40% buffer B (buffer A: 2% ACN, 0.1% FA, 0.025% TFA; buffer B: 99.9% ACN, 0.1% FA) at a 40 μL/min flow rate.

Samples were also run on a Orbitrap™ Ascend Tribrid™ mass spectrometer (Thermo Fisher) to obtain MS/MS information for both determining N-glycosylation forms and identifying pepsin digested peptides by Byonic and Byos (Protein Metrics). Peptide mass spectra were further confirmed on DriftScope (Waters) and identified with specific retention time and drift time. HDExaminer (Sierra Analytics) was used to parse the mass spectrometer datasets and calculate deuterium uptake of each peptide identified by the specific retention time and drift time.

### SDS-PAGE and western blot

For all the SDS-PAGE gels using Coomassie Blue staining, 11 μL spike samples were mixed with 1 μL DTT (0.5M) and 4 μL 4× NuPAGE™ LDS loading dye (Invitrogen) before loading to 4-12% NuPAGE™ Bis-Tris gel (Invitrogen) wells. SDS-PAGE gels were run at 120 Volts for 10 minutes and 150 Volts for 60 minutes in MES SDS running buffer (Invitrogen), followed by gel collection and PageBlue™ (Thermo Scientific) staining.

For SDS-PAGE gels proceeding to western blots, 5 μL spike samples were mixed with 1 μL DTT (0.5M) and 2 μL 4× loading dye, loaded to the SDS-PAGE gel wells, and run at 120 Volts for 80 minutes. The peptides on the gel were then transferred to the Immobilon^®^-FL PVDF membrane (Millipore) using 10% methanol supplemented NuPAGE™ transfer buffer (Invitrogen) at 30 Volts for 2 hours. All membranes were blocked in 5% non-fat milk overnight, followed by incubation in primary antibody, respectively, in room temperature for 1 hour. Primary antibody against spike S2 subunit (1A9, mouse, mAb, Invitrogen, MA5-35946) was used in 1:2000; against Strep-tag (mouse, mAb, GenScript, A01732S) was used in 1:2000; against His-tag (mouse, mAb, Invitrogen) was used in 1:2000 tris-buffered saline-Tween (TBST) solution. Secondary antibody anti-mouse IgG was used in 1:5000 5% non-fat milk blocking buffer in room temperature for 1 hour. The membranes were imaged on the Odyssey^®^ M-XS imaging system (LI-COR Biosciences) using 700 nm channel and optimized exposure intensity.

### Bio-layer interferometry

Bio-layer interferometry was used to measure binding kinetics between spikes and human ACE2 receptor or neutralizing antibodies. Octet^®^ anti-human Fc capture (AHC) biosensors (Sartorius) were pre-incubated in assay binding buffer (10 mM phosphate-buffered saline (PBS), pH 7.4, 0.1% bovine serum albumin (BSA), 0.05% Tween 20) for at least 15 minutes. Spike samples were 2-fold serial diluted with binding buffer from 200 nM to 3.1 nM. Ligand samples were also prepared with binding buffer at 2 μg/mL concentration for dimeric human ACE2-Fc and 1 μg/mL for antibodies. The binding assay was performed on an Octet^®^ BLI system (Sartorius) with 60s ligand loading, 180s association and 180s dissociation processes. BLI kinetic data were processed by ForteBio Data Analysis software (v11.0) with 1:1 binding model fitting.

### Negative-stain electron microscopy

SEC purified spike trimers were diluted to 20 ng/μL for negative-stain electron microscopy (nsEM). Carbon film 300 mesh copper grids (Electron Microscopy Sciences) were glow discharged for 30 seconds in a vacuum chamber. 3 μL of each diluted spike sample was added onto the grid (carbon side), incubated for 1 minute before blotting off with a piece of folded filter paper. The grid was then incubated in one drop of nano-W (Methylamine Tungstate, Nanoprobes) for 1 minute, followed by blotting off excess stain solution to leave a thin layer. The air-dried grid was saved in the grid box and imaged on a FEI Tecnai G2 Spirit TEM (120 kV) with a Gatan Ultrascan 4000 CCD detector. Data collection was performed using Leginon with an image pixel size of 1.6 Å.

### Statistical analysis

Statistical analysis was done for pairwise comparison between HDX-MS deuterium uptake levels. Technical replicates (n = 5 for 3-second, n = 4 for 1-minute and 15-minute time points) were prepared and run for all five constructs during HDX-MS. Two-tailed t-test was used to report the statistics. All uptake plots include error bars indicating the standard derivations from (n > 3) replicates. Error bars less than 1.5% were masked by the plotted data points on the plots.

## Notes

### Competing Interest Statement

The authors have declared no competing interest.

